# SIRT4 Controls Macrophage Function and Wound Healing through Control of Protein Itaconylation in Mice

**DOI:** 10.1101/2025.05.12.653532

**Authors:** Kristin A. Anderson, Beverly deSouza, Pol Castellano-Escuder, Zhihong Lin, Olga R. Ilkayeva, Michael J. Muehlbauer, Paul A. Grimsrud, Matthew D. Hirschey

## Abstract

Proper regulation of inflammatory responses is essential for organismal health. Dysregulation can lead to accelerated development of the diseases of aging and the aging process itself. Here, we identify a novel enzymatic activity of the mitochondrial sirtuin SIRT4 as a lysine deitaconylase that regulates macrophage inflammatory responses. Itaconate is a metabolite abundantly produced in activated macrophages. We find it forms a protein modification called lysine itaconylation. Using biochemical and proteomics approaches, we demonstrate that SIRT4 efficiently removes this modification from target proteins both *in vitro* and *in vivo*. In macrophages, elevated protein itaconylation increases upon LPS stimulation, coinciding with elevated SIRT4 expression. SIRT4-deficient macrophages exhibit significantly increased IL-1β production in response to LPS stimulation. This phenotype is intrinsic to macrophages, as demonstrated by both lentiviral over-expression and acute SIRT4 knockdown models. Mechanistically, we identify key enzymes in branched-chain amino acid (BCAA) metabolism as targets of hyperitaconylation in SIRT4-deficient macrophages. The BCKDH complex component dihydrolipoamide branched chain transacylase E2 (DBT) is hyperitaconylated and has reduced BCKDH activity in SIRT4KO macrophages. Physiologically, SIRT4-deficient mice exhibit significantly delayed wound healing, demonstrating a consequence of dysregulated macrophage function. Our data reveal a novel protein modification pathway in immune cells and establish SIRT4 as a critical regulator at the intersection of metabolism and inflammation. These findings have implications for understanding immune dysregulation in aging and metabolic disease.

## INTRODUCTION

The immune system plays a critical role in maintaining organismal health throughout the lifespan, with age-related immune dysfunction emerging as a key driver of age-associated diseases and frailty. “Inflammaging,” the chronic low-grade inflammation that characterizes aging, represents a critical intersection between innate immunity and age-related pathologies^1,2^. At the cellular level, this process is orchestrated through complex metabolic reprogramming that coordinates energy metabolism with immune function^3^.

Among the key mediators of this immunometabolic regulation, itaconate has emerged as a pivotal signaling molecule that links cellular metabolism to inflammatory control.

Itaconate is a metabolite produced primarily in activated macrophages through the decarboxylation of cis-aconitate by the enzyme immunoresponsive gene 1 (IRG1)^4^. During inflammation, the tricarboxylic acid (TCA) cycle in macrophages undergoes significant remodeling, increasing the production of itaconate, which serves as both a metabolic intermediate and a signaling molecule with potent immunoregulatory properties^5^. The importance of itaconate in immune regulation is highlighted by its ability to limit inflammatory damage by inhibiting succinate dehydrogenase, activating the Nrf2 antioxidant pathway, and modulating various inflammatory signaling networks^6^.

As a reactive carbon species^7^, itaconate can covalently modify proteins through multiple mechanisms. The best characterized is its ability to modify cysteine residues through a process called 2,3-dicarboxypropylation, which alters protein function and serves as a regulatory mechanism in inflammatory signaling^6^. However, the full spectrum of protein modifications induced by itaconate and their functional consequences remain incompletely understood.

Protein post-translational modifications (PTMs) represent a critical layer of regulation that allows cells to rapidly respond to environmental changes without requiring new protein synthesis^8^. Among the enzymes that regulate these modifications, the sirtuin family of NAD?-dependent deacylases has emerged as key mediators of metabolic adaptation^9^. The mitochondrial sirtuins (SIRT3, SIRT4, and SIRT5) remove specific acyl modifications from target proteins, typically restoring their enzymatic activity and regulating metabolic flux through various pathways^10^. While SIRT3 primarily functions as a deacetylase and SIRT5 as a desuccinylase/demalonylase, the enzymatic activity and biological function of SIRT4 have been more challenging to define.

Recent studies revealed that SIRT4 possesses unique deacylase activities, removing acyl moieties such as hydroxymethylglutaryl (HMG), methylglutaryl (MG), and glutaryl groups from lysine residues^11^. These findings established SIRT4 as a regulator of branched-chain amino acid (BCAA) metabolism, with significant implications for metabolic health. Mice lacking SIRT4 exhibit accelerated development of insulin resistance and defective adipogenesis^11,12^. Despite these advances in understanding SIRT4’s metabolic functions, significant gaps remain in our knowledge of its broader physiological roles related to aging. Specifically, dSIRT4 has been shown to influence *Drosophilia* aging, where genetic ablation leads to shorted lifespan and over-expression increases lifespan^13^. While, dysregulated inflammatory responses can accelerate age-related diseases, we realized the role of SIRT4 in immune regulation remains unexplored. Given SIRT4’s established ability to remove structurally similar acyl modifications, we hypothesized that it might target protein modifications derived from immunometabolites. Thus, we set out to investigate whether SIRT4 can remove modifications from proteins induced by itaconate, a key immunomodulatory metabolite, thereby influencing macrophage function and inflammatory responses.

## RESULTS

### SIRT4 possesses deitaconylase activity that targets a novel protein modification

We previously identified a group of chemically similar modifications that SIRT4 targets for removal. Specifically, SIRT4 removes negatively charged 5- and 6-carbon moieties called glutarylation, HMGylation, MGylation, and MGcylation^11^. These modifications to lysine residues are derived from non-enzymatic interactions with reactive acyl-CoA species. Following our discovery of these novel targets, molecular modeling using the SIRT4 crystal structure predicted that SIRT4 might more efficiently remove dimethylsuccinate as a substrate^14^. Although dimethylsuccinate is not a known physiological metabolite, its structural resemblance to itaconic acid/itaconate (Fig. 1A), a molecule abundantly produced in activated macrophages, led us to hypothesize that SIRT4 might remove protein itaconylation.

**Figure 1.**
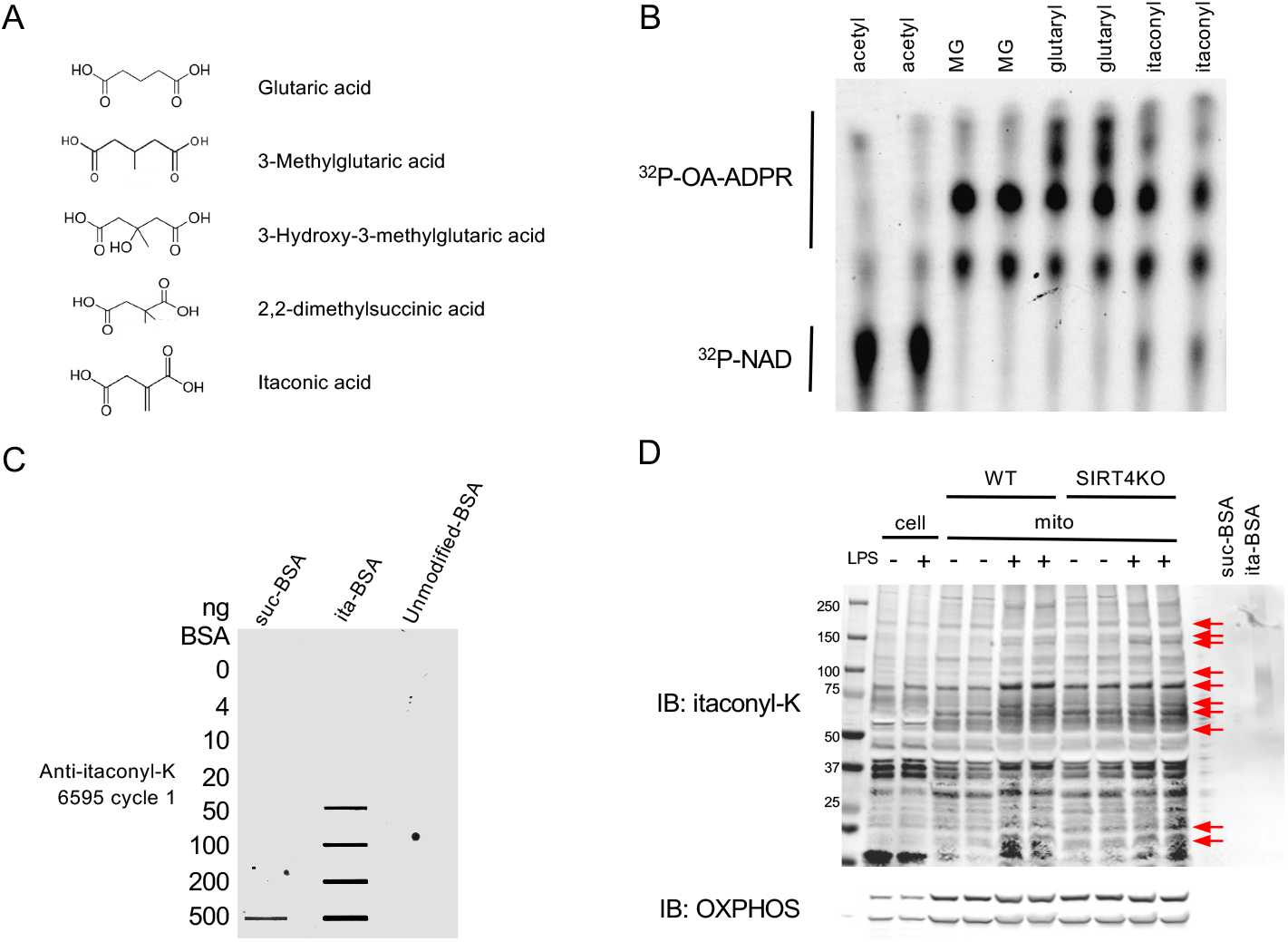
SIRT4 possesses deitaconylase activity and removes lysine itaconylation (A) Chemical structures of itaconate and other structurally similar acyl modifications targeted by SIRT4, including glutaryl, methylglutaryl (MG), and hydroxymethylglutaryl (HMG) groups. (B) SIRT4 deitaconylase activity demonstrated in vitro using NAD^+^ consumption assay. Recombinant SIRT4 was incubated with chemically itaconylated BSA in the presence of ^32^P-NAD^+^. Formation of ^32^P-O-acyl-ADPR (upper band) indicates active deacylation. (C) Validation of anti-itaconyl-lysine antibody specificity. Western blot showing antibody reactivity against unmodified BSA, succinylated BSA (suc-BSA), and itaconylated BSA (ita-BSA), demonstrating specificity for itaconyl-lysine over structurally similar succinyl-lysine. (D) Protein itaconylation increases in macrophages upon LPS stimulation. Western blot of whole-cell and mitochondrial lysates from wild-type and SIRT4KO bone marrow-derived macrophages (BMDMs) with or without LPS stimulation (10ng/ml, 24h), probed with anti-itaconyl-lysine antibody. Red arrows indicate where SIRT4KO samples show enhanced itaconylation signal at baseline, upon LPS stimulation, or both compared to wild-type in mitochondrial fractions.

To test this hypothesis, we chemically itaconylated BSA using an acid anhydride treatment and incubated it with recombinant SIRT4. We monitored the reaction for consumption of ^32^P-NAD^+^ and formation of ^32^P-O-acyl-ADPR, established indicators of SIRT4 activity. We observed robust deitaconylase activity, comparable to other known deacylase activities of SIRT4 (Figure 1B), revealing a previously unidentified enzymatic function we call lysine deitaconylation. This novel enzyme activity suggested that a corresponding protein modification could exist in mammalian cells as a physiological SIRT4 target.

While itaconate has been shown to induce cysteine modifications (2,3-dicarboxypropylation) that modulate immune responses (Mills et al., 2018), lysine itaconylation remains unexplored. As a first step toward identifying lysine itaconylation *in vivo*, we generated a polyclonal itaconyl-lysine antibody to visualize global changes in itaconylation. We used itaconylated-BSA as an antigen to establish a novel antibody (YenZyme Antibodies, LLC, San Francisco, CA). We measured itaconyl-lysine serum reactivity against unmodified BSA, succinylated BSA (suc-BSA), and itaconylated BSA (ita-BSA) (Fig. 1C) to test specificity. These data demonstrate that our antibody has a higher specificity for itaconyl-lysine than the structurally similar succinyl-lysine moiety.

To determine whether lysine itaconylation occurs under physiological conditions, we measured protein itaconylation levels in whole-cell and mitochondrial lysates from bone marrow-derived macrophages (BMDMs) isolated from wild-type and germline *Sirt4^-/-^* (KO) mice. Itaconate and cysteine 2,3-dicarboxypropylation increases upon LPS stimulation^4,6^. Thus, we predicted that lysine itaconylation might also increase. We observed increased mitochondrial protein itaconylation upon LPS stimulation in wild-type mice (Fig. 1D). Comparing WT to KO samples revealed stronger signals for specific bands in KO samples either at baseline, upon LPS stimulation, or both (Fig. 1D, arrows), consistent with the hypothesis that SIRT4 is a lysine deitaconylase and its removal leads to hyperitaconylation.

### SIRT4 regulates macrophage inflammatory responses by controlling IL-1β production

Itaconate plays a crucial role in regulating inflammation^15^ and is primarily produced in macrophages by the mitochondrial enzyme aconitate decarboxylase (*Acod1* gene, IRG1 protein)^4,5^. Given the link between itaconate and inflammation, we hypothesized that SIRT4 might regulate immune function. To characterize the immunological consequences of SIRT4 deletion, we first measured serum cytokine levels using the V-PLEX Mouse Cytokine 19-Plex Kit in wild-type (WT) and SIRT4KO mice at 2- and 7-hours after intraperitoneal injection of lipopolysaccharide (LPS). SIRT4KO mice exhibited significantly elevated levels of multiple cytokines compared to WT controls (Figure 2A). We observed a marked increase in the pro-inflammatory cytokine IL-1β (2.5-fold, p<0.001) in SIRT4KO mice 2 hours after LPS stimulation (Figures 2A,B), implying that SIRT4KO mice have exacerbated pro-inflammatory macrophages in response to LPS. Pathway analysis of the differentially expressed cytokines revealed a significant enrichment in inflammatory response pathways (Figure 2C). Specifically, inflammatory mediators (IL-15, IL-17A, IL-27p28/IL-30, IL-33, IFN-γ, IL-1β, IL-2, IL-5) and T cell-related cytokines (IL-12p70, p=0.002; IL-4, p=0.008) were significantly increased in SIRT4KO mice 2 hours after LPS stimulation. This systemic enhancement of multiple pro-inflammatory pathways suggests that SIRT4 serves as a critical negative regulator of the acute innate immune response to endotoxin challenge.

**Figure 2:**
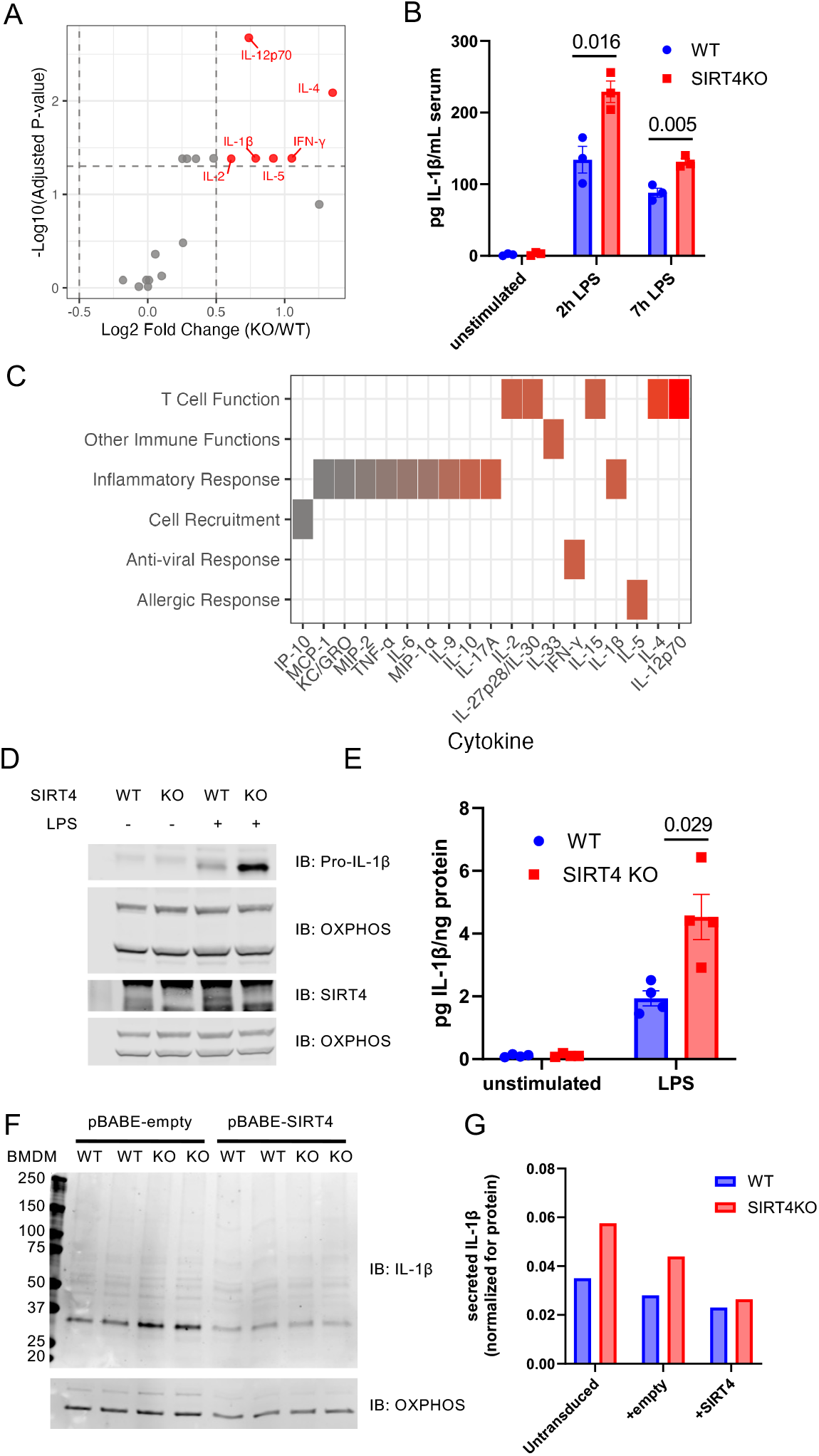
SIRT4 regulates macrophage inflammatory response by controlling IL-1β production. (A) Differential serum cytokine abundance between SIRT4KO and WT mice after intraperitoneal injection of LPS (2μg/g body weight). Serum was collected before and after LPS stimulation (2h time point shown) and analyzed for inflammatory cytokines using V-PLEX Mouse Cytokine 19-Plex Kit. Each point represents a cytokine, with the x-axis showing the log2 fold change (KO/WT) and the y-axis the -log10 of the adjusted p-value (limma). Cytokines that are significantly different between genotypes (adjusted p-value < 0.05 and absolute log2 fold change > 0.5, dotted lines) are highlighted in red. (B) Elevated serum IL-1β in SIRT4KO mice after intraperitoneal injection of LPS (2μg/g body weight). Serum was collected before and after LPS stimulation and analyzed for inflammatory cytokines using V-PLEX Mouse Cytokine 19-Plex Kit. Data are shown as mean ± SEM; *p<0.05. (C) Cytokine Functional Classification Heatmap. Cytokines grouped by immune function, with color intensity (p-values ranging from 0.002 to 0.967; red to gray) indicating statistical significance between KO and wild-type mice. Inflammatory mediators and T cell-related cytokines show the strongest differential abundance. (D) Western blotting of wild-type and SIRT4KO BMDM cell extracts showing elevated pro-IL-1β in KO samples (upper panel) and increased SIRT4 expression in WT samples following LPS stimulation (middle panel). OxPhos was used as a loading control (lower panel). (E) Increased IL-1β secretion from SIRT4KO BMDMs after LPS stimulation. BMDMs from wild-type and SIRT4KO mice were stimulated with LPS (10ng/ml) for 24h, and secreted IL-1β was quantified by ELISA and normalized to total protein. Data are shown as mean ± SEM; *p<0.05 (F & G) Macrophage-intrinsic SIRT4 directly controls inflammatory responses. BMDMs from WT and SIRT4KO mice were transduced with either empty pBABE vector (empty) or SIRT4-expressing pBABE vector (SIRT4) using a retroviral delivery system, then stimulated with LPS. Pro-IL-1β protein levels (Western blot, F) and secreted IL-1β (ELISA, G) demonstrate rescue of inflammatory phenotype upon SIRT4 reintroduction.

To determine whether macrophages were responsible for this phenotype, we generated bone marrow-derived macrophages (BMDMs) from WT and SIRT4KO mice and analyzed cytokine production after LPS stimulation. Consistent with our *in vivo* observations, BMDMs from KO mice exhibited significantly higher pro-IL-1β protein levels before and after LPS stimulation (Fig. 2D), as well as increased levels of secreted IL-1β (Fig. 2E). Intriguingly, we also observed SIRT4 is constitutively expressed in BMDMs and its levels increase upon LPS stimulation (Fig. 2D, middle panel), suggesting a cell-intrinsic role in regulating macrophage activation and inflammatory responses. Altogether, these data support the model that SIRT4 negatively regulates IL-1β in macrophages and its absence leads to elevated production in response to inflammatory stimuli.

### Macrophage-intrinsic SIRT4 directly controls inflammatory responses

To test a direct role of SIRT4 within macrophages in response to LPS, we performed complementary gain- and loss-of-function experiments. First, we ectopically expressed SIRT4 in BMDMs from both SIRT4 WT and KO mice using a lentiviral system, then stimulated the cells with LPS. SIRT4 expression in KO BMDMs restored both pro-IL-1β (Fig. 2F) and secreted IL-1β (Fig. 2G) to levels comparable to those observed in WT BMDMs, confirming that SIRT4 expression in macrophages is sufficient to normalize inflammatory responses. These data indicate a direct role for SIRT4 in regulating the response to LPS. To further validate these findings, we knocked down endogenous SIRT4 in the monocyte/macrophage-like RAW cell line using siRNA (Supplemental Fig. 1). Three days after transfection, LPS stimulation resulted in significantly increased IL-1β production relative to control cells (Supplemental Fig. 1), phenocopying the response observed in BMDMs from SIRT4KO mice. These complementary approaches demonstrate that modulating SIRT4 levels within macrophages directly affects their inflammatory response to LPS.

### SIRT4 regulates itaconylation of BCAA catabolic enzymes in macrophages

To investigate SIRT4’s role in macrophage activation, we performed label-free mass spectrometry-based proteomics on LPS-stimulated bone marrow-derived macrophages (BMDMs) from wild-type and SIRT4KO mice. LPS stimulation caused significant protein abundance changes at the 24h time point, consistent with transition from basal to activated state^16^. Importantly, aside from Sirt4 protein, we detected no significant differences in overall protein abundances between WT and KO BMDMs (ProteomeExchange PXD063862). Next, we examined protein itaconylation sites by re-analyzing our samples using a variable modification search with a monoisotopic mass of 112.0160 Da added to lysine residues (Fig. 3A). From 47,966 peptides identified at 1% global false discovery rate (FDR), 158 peptides contained itaconylated lysine. Most proteins (97.3%) contain only a single specific itaconylation site, with 2.0% containing two sites and 0.7% having three or more sites (Fig. 3B). Because the analysis used unenriched samples due to technical limitations preventing itaconylated peptide enrichment, we consider this data a preliminary survey of the itaconylation landscape.

**Figure 3:**
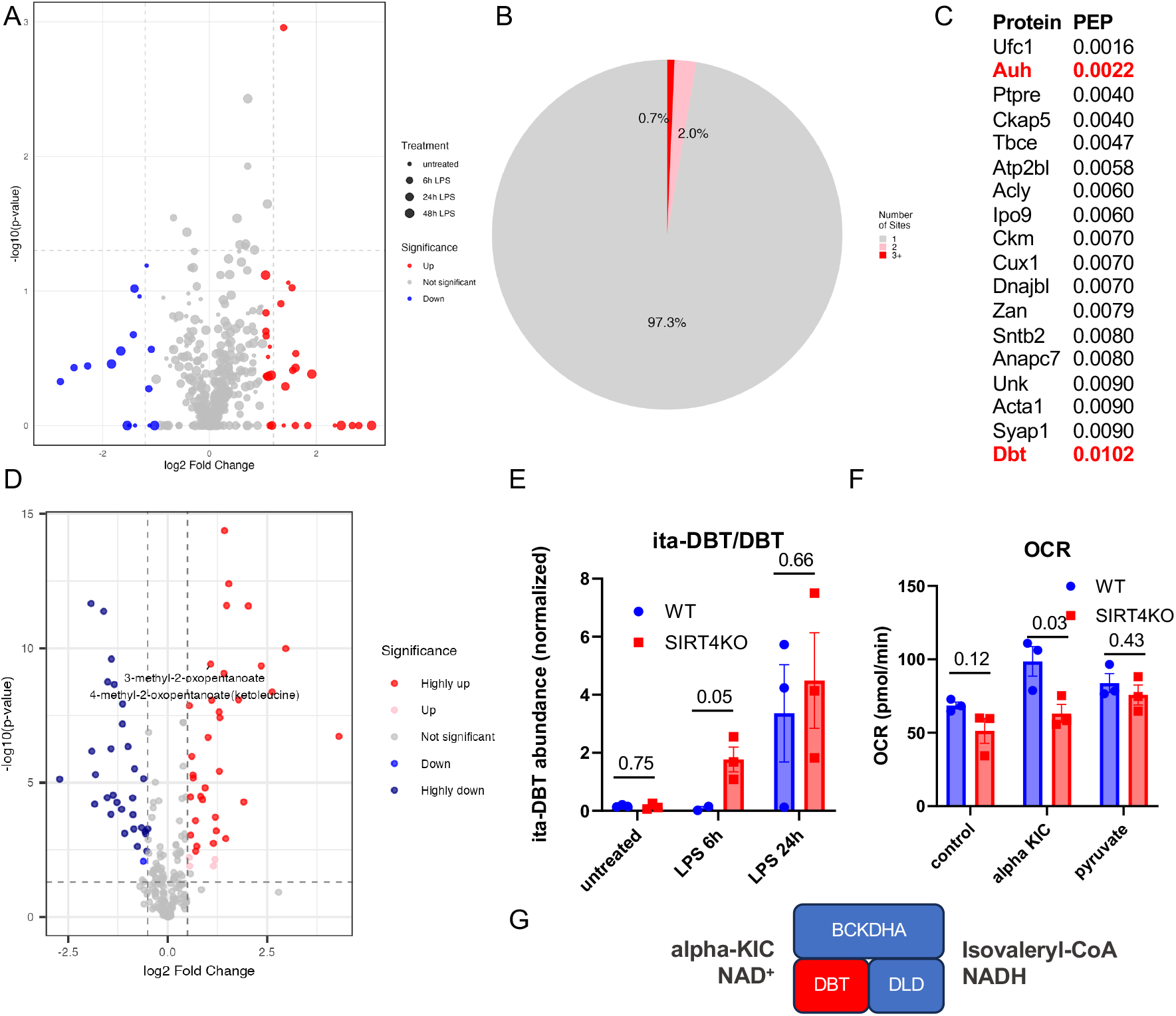
SIRT4 regulates BCAA metabolism in macrophages through control of protein itaconylation. (A) Differential protein itaconylation changes in BMDMs in SIRT4KO mice compared to wild-type following LPS stimulation for 0h, 6h and 24h. Log2 fold change (KO/WT x-axis) vs. statistical significance (−log10 p-value, y-axis). Red: upregulated sites (p<0.05, log2FC>0.5); blue: downregulated sites (p<0.05, log2FC<-0.5); gray: non-significant changes. Dashed lines indicate significance threshold (p=0.05) and fold change thresholds (±0.5). (B) Itaconylation Sites Per Protein. Distribution of proteins based on number of distinct itaconylation sites. Single site: 97.3%; two sites: 2.0%; three or more sites: 0.7%. Most proteins contain only a single specific modification site. (C) List of high-confidence itaconylated peptides identified by mass spectrometry, with posterior error probability (PEP) scores ≤0.010 (1% local FDR). Proteins involved in BCAA metabolism (AUH and DBT) are highlighted in red. (C) Untargeted metabolomics from BMDMs in SIRT4KO mice compared to wild-type following LPS stimulation for 0h, 6h and 24h. The x-axis represents log2 fold change, with negative values indicating downregulation and positive values indicating upregulation. The y-axis shows -log10(p-value), where higher values indicate greater statistical significance. The plot highlights a notable metabolite: 4-methyl-5-oxopentanoate (increased) involved in BCAA metabolism. Horizontal and vertical dashed lines represent significance and fold change thresholds, respectively. (D) Analysis of itaconylated-dihydrolipoamide branched chain transacylase E2 (ita-DBT) peptide abundances indicates hyperitaconylation in LPS-stimulated BMDMs from SIRT4KO mice compared to wild-type at 6h and 24h post-stimulation. Data are shown as normalized peptide abundance from mass spectrometry analysis. P-values indicated from mass spectrometry analyses. (E) BCKDH complex activity is reduced in BMDMs from SIRT4KO mice. Oxygen consumption rate (OCR) was measured in permeabilized BMDMs using the XF24 Seahorse analyzer. NADH produced from either NAD^+^ alone (untreated, as negative control), α-ketoisocaproate (αKIC, as a substrate to test BCKDH), or pyruvate (as a test of PDH). Data are normalized to protein content and shown as mean ± SEM; *p<0.05; n=3 independent experiments. (F) Schematic showing DBT in the BCKDH complex

As the fraction of itaconylated peptides identified at 1% global FDR (Fig. 3C-D) was low (0.3%) we set a conservative posterior error probability (PEP) score threshold at 0.010 for validation of individual itaconyl peptide hits (Figure 3C), with several proteins containing itaconylated peptides with PEP ≤ 0.010, or 1.0% local FDR^17^.

To better understand the pathways that could be controlled by SIRT4 in the macrophage, we performed untargeted metabolomics on LPS-stimulated BMDMs from SIRT4KO mice compared to wild-type, again at 0-, 6-, and 24-hours of LPS stimulation. This analysis revealed changes in several metabolites that were significantly altered (Fig. 3D). Next, we performed multi-omics integration of proteomic, itaconyl-proteomic, metabolomic, and inflammatory datasets using an algorithm we recently developed called GAUDI^18^. Analyses revealed distinct clustering patterns integrating both genotype and LPS treatment time points (Supplemental Fig. 2). Standard UMAP visualization demonstrated clear separation between samples, with LPS treatment duration (0h, 6h, and 24h) forming distinct clusters (Supplemental Fig. 2B). Feature importance analysis identified several known contributors to macrophage function, including TRAP1 (TNF Receptor Associated Protein 1) and IFNGR1, as top contributors to UMAP1 and UMAP2 variance (Supplemental Figs. 2C and 2D). Gene Ontology enrichment analysis of the integrated dataset revealed significant enrichment of biological processes related to hydrogen peroxide signaling and programmed cell death (Supplemental Fig. 2E), with the most significant terms involving key macrophage LPS-response pathways. This approach identified molecular features known to contribute to macrophage activation^18^, demonstrating that integration of metabolomic, proteomic, and itaconylated peptide datasets provides comprehensive insights into the molecular changes occurring during this perturbation.

However, the signals from LPS treatment masked any signals from the SIRT4 ablatoin genotype (Supplemental Fig. 2A), indicating that we needed to interrogate specific genotype differences within each time point. Returning to our high confidence itaconylated protein targets, we observed two mitochondrial proteins, AUH and DBT, participating in branched-chain amino acid (BCAA) oxidation (Fig. 3C), a pathway previously shown to be regulated by SIRT4^11,12^. Interestingly, AUH participates in both BCAA oxidation and itaconyl-CoA oxidation pathways^19^. DBT is the E2 lipoamide acyltransferase component of the branched-chain alpha-keto dehydrogenase (BCKDH) complex that catalyzes the key step in BCAA oxidation. AUH itaconyl-peptide abundances showed no significant differences between SIRT4 WT and KO BMDMs.

We detected significantly higher DBT itaconyl-peptides in LPS-stimulated BMDMs from SIRT4KO mice (Fig. 3E). This hyperitaconylation was most pronounced at the 6h LPS time point and continued at the 24h time point, suggesting DBT might be a direct target of SIRT4’s deitaconylase activity. Furthermore, the metabolomics data show that 4-methyl-5-oxopentanoate (aka alpha-ketoisocaproate), the metabolic substrate of BCKDH, was elevated in SIRT4KO mouse BMDMs stimulated with LPS, further supporting a connection between SIRT4 and BCAA metabolism in macrophages.

Thus, we directly tested whether SIRT4 affects BCAA metabolism in macrophages by measuring BCKDH activity directly in BMDMs from WT and SIRT4KO mice. We leveraged an assay previously used by our group using the XF24 Seahorse instrument^11^. Macrophages attached to a Seahorse microplate were permeabilized, and their growth medium was replaced with defined medium containing BCKDH substrates (alpha-ketoisocaproate [αKIC], thiamine pyrophosphate [TPP], and NAD^+^). BCKDH-produced NADH feeds into the electron transport chain, making oxygen consumption rate (OCR) a readout for BCKDH activity. We measured OCR in BMDMs from WT and SIRT4KO mice using αKIC/TPP/NAD^+^ as substrate, with pyruvate/NAD^+^ as positive control and NAD^+^ alone as negative control. BMDMs from SIRT4KO mice exhibited OCR values similar to those from WT animals when supplied with either NAD^+^ alone or pyruvate/NAD^+^ (Fig. 3F). In contrast, SIRT4KO BMDMs showed significantly reduced OCR values compared to WT cells when provided with αKIC/TPP/NAD^+^ (p<0.05).

These results suggest SIRT4 influences BCAA oxidation in macrophages by promoting BCKDH activity and its absence leads to less BCAA catabolism. Altogether, these results lead to a model where SIRT4 regulates BCAA metabolism in macrophages through protein itaconylation control, by preventing excessive DBT itaconylation (Fig. 3G). This finding establishes a novel mechanistic link between SIRT4, protein itaconylation, and macrophage inflammatory response regulation through BCAA metabolism modulation.

### SIRT4 deficiency impairs wound healing *in vivo*

To investigate the physiological consequences of enhanced IL-1β production in SIRT4-deficient macrophages, we examined wound healing, because elevated macrophage activation and increased IL-1β secretion have been linked to delayed tissue repair in multiple models^20^. We assessed wound healing *in vivo* using an excisional wound model in which small skin wounds were created using a biopsy punch, and wound closure was monitored over the course of one week in both male and female mice.

Representative images from a matched pair of male WT and SIRT4KO mice show delayed wound healing in the absence of SIRT4 (Fig. 4A). In both male and female SIRT4KO mice, we observed statistically significant delays in wound closure compared to WT controls (Figs 4B-4C). This impaired healing response demonstrates a physiological consequence of the dysregulated immune function in SIRT4KO mice and establishes a link between SIRT4’s molecular function in regulating macrophage activation and its role in tissue repair processes. Overall, these findings lead to a new model for SIRT4 in regulating macrophage function and wound healing (Fig. 4D). During inflammatory activation, macrophages produce itaconate via IRG1/Acod1, leading to protein itaconylation. SIRT4 removes itaconyl modifications from key metabolic enzymes including DBT, maintaining BCKDH activity and proper BCAA catabolism. This metabolic regulation limits IL-1β production and promotes timely resolution of inflammation and wound healing. In SIRT4’s absence, hyperitaconylation of metabolic enzymes leads to dysregulated BCAA metabolism, elevated IL-1β production, and impaired wound healing.

**Figure 4:**
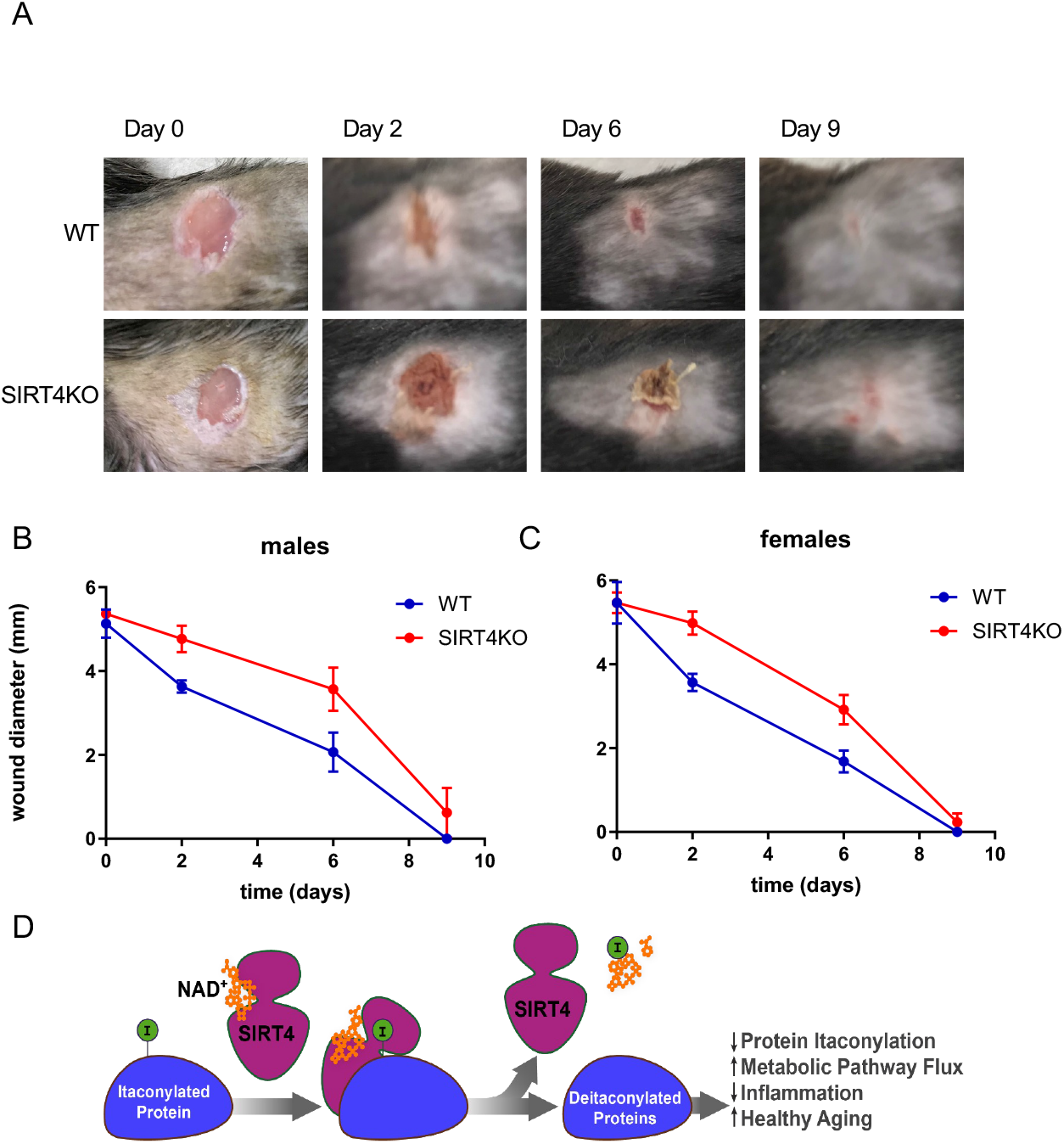
SIRT4 deficiency impairs wound healing in vivo. (A) Representative images of excisional wounds in wild-type and SIRT4KO male mice at days 0, 3, 6, and 9 post-wounding. Full-thickness dorsal wounds were created using 5mm biopsy punches. (B-C) Quantification of wound closure in male (B) and female (C) mice. Wound diameters were measured using digital calipers over 9 days in wild-type and SIRT4KO mice (n=3 per genotype). Data are shown as mean ± SEM; *p<0.05 by t-test at individual time points; ANOVA revealed statistically significant difference between genotypes (p<0.05). (D) Proposed model illustrating SIRT4’s role in regulating macrophage function and wound healing.

## DISCUSSION

The discovery of SIRT4’s deitaconylase activity adds a new dimension to the sirtuin family’s expanding repertoire of protein targets. While SIRT3 and SIRT5 have been well characterized as deacetylase and deacylase enzymes, SIRT4’s enzymatic functions have remained more elusive. This study demonstrates that SIRT4 efficiently removes itaconyl modifications from lysine residues, representing a novel enzymatic activity in immune cell function.

SIRT4 upregulation during macrophage activation occurs at a noteworthy time. We observed increased SIRT4 expression following LPS stimulation, coinciding with itaconate production’s peak^21^. This temporal relationship suggests that SIRT4 may function as a negative regulator of immune responses by limiting protein itaconylation’s duration and extent. SIRT4 likely contributes to inflammation’s resolution phase, preventing excessive inflammatory damage after macrophage activation.

Lysine itaconylation represents a novel protein modification pathway that may have evolved specifically for immune cell function regulation. Unlike other acyl modifications derived from central metabolic pathways common to most cell types, itaconate is primarily produced in activated macrophages through IRG1/Acod1^ref 5^. This restricted pattern suggests itaconylation serves specialized regulatory roles in immune responses. Our findings extend the paradigm established by previous studies on cysteine itaconylation by demonstrating that lysines are also subject to itaconylation^6^, potentially affecting a broader range of protein functions.

Our proteomics analysis revealed that BCAA metabolism enzymes, particularly DBT (a BCKDH complex component), are hyperitaconylated in SIRT4KO macrophages. This observation, coupled with reduced BCKDH activity in SIRT4KO BMDMs, establishes a mechanistic link between SIRT4’s enzymatic activity and BCAA catabolism regulation in immune cells. These observations align with previous work demonstrating SIRT4’s role in controlling BCAA oxidation in liver and adipose tissue^11,12^, and extend these findings to the immune compartment.

The potent effect on IL-1β production, rather than broad dysregulation of all inflammatory cytokines, is particularly intriguing. Recent studies have revealed that BCAA metabolism plays a crucial role in determining macrophage polarization and cytokine production. The increase in IL-1β observed in our SIRT4KO model suggests specific effects on macrophage activation pathways rather than on global immunologic alterations.

Further, our findings suggest that SIRT4 may influence this process by maintaining BCKDH activity through deacylation, thereby preventing BCAA accumulation and limiting IL-1β production. This specific regulatory effect may reflect the integration of BCAA metabolism with inflammasome activation. Previous studies have shown that metabolic perturbations can selectively influence IL-1β processing through NLRP3 inflammasome regulation^22^. The connection between BCAA metabolism and IL-1β production represents an important link between cellular metabolism and inflammatory control^23,24^, with SIRT4 potentially serving as a key mediator at this interface.

The delayed wound healing observed in SIRT4KO mice demonstrates a physiological consequence of dysregulated macrophage function and establishes a connection between molecular mechanisms and tissue-level outcomes. Macrophages are pivotal in wound healing, initiating the response and aiding in closure and repair. The impaired wound healing phenotype in our SIRT4KO model provides compelling evidence that the molecular alterations we observed have significant physiological consequences.

Supporting this idea, macrophage-secreted IL-1β plays a crucial role in skin inflammation and wound healing dynamics. Elevated IL-1β maintains a pro-inflammatory state that can impede healing. Studies have shown that blocking IL-1β shifts macrophages toward a healing phenotype, enhancing repair in diabetic wounds. The increased IL-1β production in SIRT4-deficient macrophages likely contributes directly to the delayed wound healing phenotype we observed.

This mechanism is particularly relevant in the context of “inflammaging,” as elevated IL-1β in aging has been shown to impair healing by increasing inflammation in skin wounds. Given that SIRT4 overexpression extends lifespan in *Drosophila*, our findings provide a potential mechanistic link between SIRT4 activity, inflammation, and aging. By regulating macrophage activation through control of protein itaconylation and BCAA metabolism, SIRT4 may contribute to organismal health and longevity by limiting chronic inflammation and promoting efficient tissue repair.

The wound healing defect in SIRT4KO mice also parallels observations in metabolic diseases such as diabetes, where impaired macrophage function and elevated IL-1β contribute to poor tissue repair. Previous studies have demonstrated that SIRT4 deficiency leads to accelerated insulin resistance and glucose intolerance, conditions associated with impaired wound healing. This suggests that the metabolic and immune dysfunctions observed in SIRT4 deficiency may be mechanistically linked through shared pathways, positioning SIRT4 as a potential therapeutic target for conditions characterized by both metabolic dysfunction and impaired tissue repair.

In conclusion, our study identifies SIRT4 as a lysine deitaconylase that regulates macrophage function by controlling protein modifications on key metabolic enzymes. This mechanism links SIRT4’s established role in metabolic regulation to a novel function in immune control and tissue repair. By uncovering this previously unrecognized pathway, our findings provide new insights into the molecular mechanisms underlying immunometabolic regulation and suggest potential therapeutic targets for inflammatory conditions and age-related diseases. Because the immune system and inflammation critically influence key aspects of health and disease, our results position SIRT4 as part of a cellular program to control immuno-metabolism during inflammatory states and aging. The identification of SIRT4 as a regulator of both metabolism and inflammation through its deitaconylase activity represents a significant advance in our understanding of the molecular mechanisms that coordinate these essential cellular processes. Future studies will be needed to fully characterize the itaconylation landscape in various immune cell types and disease states, and to explore the therapeutic potential of targeting the SIRT4-itaconylation axis in inflammatory conditions and age-related pathologies.

## Supporting information

Reagents

Uncropped blots

## ACKNOWLEDGMENTS

We would like to thank Nancie McIver for assistance with immune cell profiling, Greg Taylor for assistance with animal phenotyping, Jason Locasale for assistance with nontargeted metabololmics analyses, and the entire Hirschey lab for thoughtful feedback. We would like to acknowledge funding support from the Glenn Foundation (MDH), the Duke Cancer Institute P30 Cancer Center Support Grant P30CA014236 (MDH), from the National Institutes of Health and the NIA R01AG045351 (MDH) and 1R21AG080334 (MDH).

## SUPPLEMENTAL FIGURE LEGENDS

**Supplemental Figure 1.** RAW264.7 cells transfected with SIRT4 siRNA or negative control siRNA for 3 days were stimulated with LPS (500ng/ml) for 24h. IL-1β production was measured by Western Blotting.

**Supplemental Figure 2. Multi-omic integration of macrophage responses to LPS treatment using GAUDI.** A-B. UMAP visualization of integrated metabolomic, proteomic, and itaconylated peptide data, with points representing individual samples colored by genotype (WT vs. KO) (A) or LPS treatment duration (0, 6, or 24 hours) (B). The UMAP space reveals that LPS treatment duration dominates the clustering pattern, with distinct grouping of LPS_0 samples (right clusters) separating from LPS_6 and LPS_24 samples (left clusters). C-D. Top 10 features contributing to UMAP1 (C) and UMAP2 (D) dimensions based on Gini index values from the Random Forest model.

Features are color-coded by source (blue: metabolites, orange: proteins). E. Gene Ontology enrichment analysis of features contributing to the integrated GAUDI model shows significant enrichment of hydrogen peroxide-mediated programmed cell death pathways, mitochondrial depolarization processes, and ubiquitin ligase complex activity. These biological processes align with known LPS-induced inflammatory responses and oxidative stress pathways in macrophages, explaining the dominant effect of LPS treatment observed in the clustering patterns.

**Supplemental Table 1.** Proteomics Analysis of Protein Itaconylation in Macrophages. This table presents mass spectrometry analysis of itaconylated proteins in wild-type and SIRT4KO bone marrow-derived macrophages with and without LPS stimulation. Tab 1: Proteins - Master list of all identified proteins with accession numbers and confidence scores. Tab 2: Peptide Groupus_Itaconyl - Detailed listing of itaconylated peptides showing modified lysine residues. Tab 3: Itaconyl_Perseus Output - Quantitative abundance values for itaconylated peptides across experimental conditions. Tab 4: Itaconyl Summary - Statistical analysis of differentially itaconylated proteins between genotypes, highlighting metabolic enzymes including Dbt and Auh. The analysis identified 158 unique itaconylated peptides with most proteins containing single modification sites, revealing significant differences between SIRT4KO and wild-type macrophages after LPS stimulation. Raw data are available through ProteomeXchange (accession number PXD063862) and jPOST repository (accession number JPST003806).

## Declaration of generative AI and AI-assisted technologies

During the preparation of this work the authors used quartzreport and carbondraft, AI-assisted tools by Heureka Labs, to create code to analyze data and enhance the readability of the text. After using these tools, the authors reviewed and edited the content and take full responsibility for the publication’s content.

## Notes

https://zenodo.org/records/15372985

### Competing Interest Statement

The authors have declared no competing interest.

