## Supplementary figures and images for "SIRT4 Controls Macrophage Function and Wound Healing through Control of Protein Itaconylation in Mice"

### Supplemental Figures

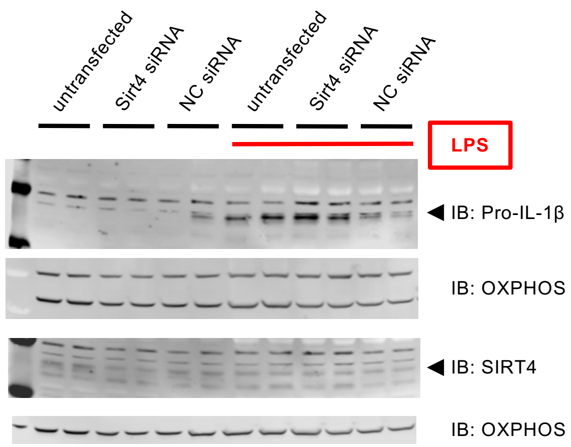


**Supplemental Figure 1 (Corresponds to Fig. 2)**

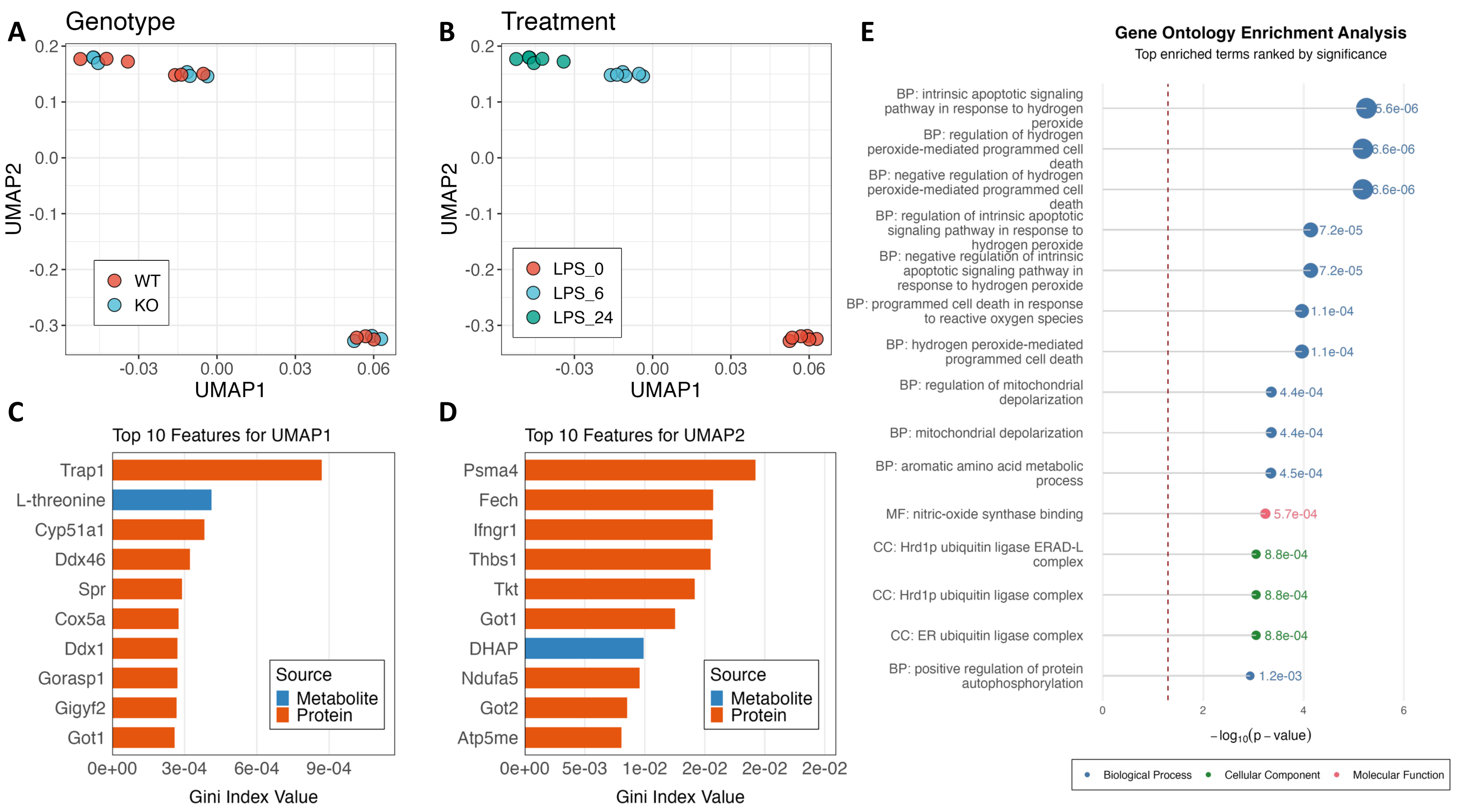


**Supplemental Figure 2 (Corresponds to Fig. 3)**
