## Supplementary material for "SIRT4 Controls Macrophage Function and Wound Healing through Control of Protein Itaconylation in Mice": Methods

**STAR METHODS**

EXPERIMENTAL MODEL AND SUBJECT DETAILS

Mice

Whole-body SIRT4KO mice were obtained from the Jackson Laboratory (Bar Harbor, ME, stock #012756) and backcrossed for 7 generations onto a C56BL/6J background obtained from Jackson Laboratory (Bar Harbor, ME, stock #000664). These mice were then backcrossed for another 3 generations onto the C57BL/6NJ background obtained from the Jackson Laboratory (Bar Harbor, ME, stock #005304) to re-introduce a functional nucleotide transhydrogenase (Nnt) gene, which is missing in the commonly used C57BL/6J mice. Sirt4tm1a(EUCOMM)Hmgu mice were obtained from the International Mouse Phenotyping Consortium. These mice were crossed with mice expressing Flp recombinase (B6.129S4-Gt(ROSA)26Sortm2(FLP*) (Sor/J, stock no. 012930; The Jackson Laboratory) to generate SIRT4flox/flox mice. SIRT4flox/flox mice were crossed to LysM-cre mice (Jackson Laboratory #004781) that express the Cre recombinase under the mouse LysM promoter, to generate macrophage-specific SIRT4KO mice (hereafter referred to as LysM-cre SIRT4KO mice). All mice were group-housed on a 12h light/dark cycle with free access to water and PicoLab Diet 20 (LabDiet #5053, St. Louis, MO). Both male and female mice aged 8-12 weeks were used for all experiments. All animal procedures were conducted in accordance with guidelines approved by the Duke University Institutional Animal Care and Use Committee.

Cell lines

L929 cells (ATCC, CCL-1) were cultured in DMEM supplemented with 10% FBS and 1X Penicillin/Streptomycin. The medium was collected after 2-3 days of culture with confluent cells, filtered through a 0.2-micron filter, and used for BMDM differentiation. RAW 264.7 cells (ATCC, TIB-71) were maintained in DMEM supplemented with 10% FBS and 1X Penicillin/Streptomycin. 293T cells (ATCC, CRL-3216) were used for retrovirus production and maintained in DMEM with 10% FBS. BL21 (DE3) pLysS bacteria were used for recombinant protein expression and were cultured in LB (Miller's) broth containing 100μg/mL ampicillin. All cells were maintained at 37°C in a humidified incubator with 5% CO2 unless otherwise specified.

Primary cell isolation and culture

Bone marrow-derived macrophages (BMDMs) were generated from bone marrow isolated from femurs and tibias of 8-12 week old mice. Both male and female mice were used for bone marrow isolation. Following isolation, bone marrow cells were cultured in macrophage differentiation medium (DMEM, 10% FBS, 30% L929 cell conditioned medium, 1X Pen/Strep) for 7 days to generate mature BMDMs. Sex of mice used for each experiment is specified in the figure legends.

METHOD DETAILS

Mouse genotyping

Whole-body SIRT4KO mice genotypes were determined using the Sirt4 F, Sirt4 R1, and Sirt4 R2 primers (sequences provided in Key Resources Table). LysM-Cre SIRT4KO mice genotyping was determined using the oIMR3066: 5'-CCC AGA AAT GCC AGA TTA CG-3' (Mutant primer), oIMR3067: 5'-CTT GGG CTG CCA GAA TTT CTC-3' (Common primer), and oIMR3068: 5'-TTA CAG TCG GCC AGG CTG AC-3' (Wildtype primer).

Chemical acylation of BSA

Bovine serum albumin (BSA), fatty acid free (Sigma #A7030) at 2mg/mL in 0.1M sodium bicarbonate, pH 8.0, was mixed with acetic, glutaric, 3-methyl-glutaric, or itaconic anhydride (Sigma #259926) at a 10-fold molar excess over BSA lysine residues to form acetyl, glutaryl, 3-methyl-glutaryl, and itaconyl-BSA, respectively. The reactions were incubated 30min at room temperature with continuous mixing, following which 20mM Tris HCl, pH 8.0 was added to neutralize unreacted anhydride. BSA lysine modification was verified by gel electrophoresis and by mass spectrometry analysis by the Duke DMPI Metabolomics/Proteomics Core.

SIRT4 in vitro activity assay

As described previously (Anderson et al., 2017), reactions were performed in a total volume of 10μl in a reaction mix containing 50mM Tris HCl, pH 8.0, 4mM MgCl2, 50mM NaCl, 0.5mM DTT, and 0.5μCi 32P-nicotinamide adenine nucleotide (NAD) (Perkin Elmer #NEG023, 800Ci/mmol) in the presence of 2μg acylated BSA and 1-2μg recombinant SIRT4 protein. After a 1h incubation at 37°C, 1.5μl of each reaction was spotted onto a silica-gel coated thin layer chromatography plate (Sigma-Aldrich #60768) and eluted with ethanol:aqueous ammonium bicarbonate (1M) at a ratio of 7:3. The plates were air dried and exposed to a Kodak storage phosphorimager screen (GE Healthcare #SD230) and the signal detected using a STORM820 phosphorimager (GE Healthcare).

Antibodies and Western blotting

Commercial antibodies used were as follows: anti-IRG1 (Cell Signaling Technology, #77510), anti-OxPhos (abcam, ab110413), and anti-SIRT4 (Sigma, HPA029691). The anti-itaconyl lysine polyclonal antibody was generated by YenZym Antibodies LLC (San Francisco, CA) using the "regular rabbit antibody service". The immunogen used was itaconylated BSA prepared as described in the "Chemical acylation of BSA" section. For Western blotting analysis using LI-COR, whole-cell or mitochondrial enriched protein extracts were resolved by SDS-PAGE and transferred to nitrocellulose membranes. The membranes were blocked in LI-COR buffer (0.6X PBS, 0.25% fish gelatin; Sigma-Aldrich #G7041, 0.05% casein; Sigma-Aldrich #C3400, and 0.02% azide) for 1h at room temperature and then probed with primary antibody in LI-COR buffer/tween (LI-COR buffer with 0.1% Tween 20). After incubation with infrared dye-conjugated secondary antibodies, the blots were scanned using the LI-COR Odyssey Infrared Imaging System.

Isolation of mitochondrial enriched protein extracts from BMDM

BMDM cell pellets were homogenized in 10 volumes of STE buffer (0.25M sucrose, 10mM Tris HCl, pH 8.0, 1mM EDTA, HALT protease inhibitors; Thermo Fisher Scientific, #78420B) with 20 strokes in a chilled glass-teflon homogenizer. The homogenate was centrifuged at 700 x g for 10min at 4°C and the resulting supernatant centrifuged at 7,000 x g for 10min at 4°C to produce a mitochondria-enriched pellet. The mitochondrial-enriched pellet was solubilized in RIPA buffer for Western blotting.

Recombinant GST-SIRT4 protein expression construct

Recombinant mouse GST-SIRT4 protein was expressed from a cDNA protein expression construct encoding a truncated version of SIRT4 (residues 24-333) cloned into pGEX-6P1 glutathione S-transferase (GST) expression vector (GE Healthcare, #28-9546-48) as described in detail previously (Anderson et al., 2017).

Bacterial cell culture conditions for recombinant SIRT4 protein

A starter culture (100mL) of BL21 (DE3) pLysS bacteria transformed with SIRT4/pGEX-6P1 cDNA, was grown in LB (Miller's) broth containing 100μg/mL ampicillin overnight at 37°C. The next day 300-400mL LB was seeded with the starter culture at an OD600 of 0.05 and incubated at 37°C on an orbital shaker until the OD600 reached 0.6-1. The culture was then cooled to room temperature and induced with 0.5mM IPTG for 16h at room temperature in an orbital shaker. Cells were harvested by centrifugation (7,700 x g for 10min at 4°C) and re-suspended on ice in 15mL PBS, pH 7.4. Cell pellets were stored at -80°C until needed for subsequent protein purification.

SIRT4 purification with GST tag cleaved

Protein purification was performed according to the exact conditions described elsewhere (Anderson et al., 2017).

Isolation of mouse bone marrow and generation of bone marrow derived macrophages

Both femurs and tibia were dissected from the mouse, the muscle removed, both ends clipped from each bone, and the bones placed in DMEM containing penicillin/streptomycin (DMEM/Pen/Strep). Bone marrow was flushed from the bones with 1mL DMEM/Pen/Strep using a 27-gauge needle attached to a 1mL syringe. After breaking up the marrow by moving it up and down through the syringe the marrow was diluted to 10mL in DMEM/Pen/Strep, and the cells pelleted by centrifugation at 1,000 x g for 5min. The resulting cell pellet was loosened by flicking the tube and the red blood cells lysed by adding 1mL ACK lysis buffer for 30sec. Cells were then diluted to 6mL with DMEM/Pen/Strep and pelleted by centrifugation at 1,000 x g for 5min. Bone marrow cells were resuspended in macrophage differentiation medium (DMEM, 10% FBS, 30% L929 cell conditioned medium, 1X Pen/Strep) and plated in 10cm uncoated petri dishes (5-6 dishes per mouse). After 5 days the medium was replaced with fresh macrophage differentiation medium. Two days later differentiation medium was removed and the cells stimulated with 100ng/mL LPS (Invitrogen REF co-4976-93, stock is 2.5 mg/ml) in activation medium (DMEM, 10% FBS, 1X Pen/Strep). L929 cell conditioned medium is medium (DMEM, 10% FBS, 1X Pen/Strep) that was in contact with confluent L929 cells for 2-3 days, at which point the medium was removed and frozen at -20°C. Thawed L929 conditioned media was passed through a 0.2-micron filter before use with bone marrow cells.

IL-1β quantification

IL-1β secreted into the growth medium by BMDM or RAW cells was quantified using the mouse IL-1 beta/IL-1F2 DuoSet ELISA kit from R&D Systems (#DY401-05) and a growth medium sample size of 100-200μL per 96-well. The in vivo production of pro-inflammatory cytokines in SIRT4KO mice following an intraperitoneal injection of LPS at 2μg/g body weight, was determined using the V-Plex Mouse Cytokine 19-plex Kit (Meso Scale Diagnostics, LLC, #K15255D).

Ectopic expression of SIRT4 in BMDM

The retroviral vector pBABE with a mouse SIRT4 coding region insert or without insert (empty vector control) was used to generate retrovirus. To generate retrovirus, 100cm dishes that were 75% confluent with 293T cells were transfected with the pBABE constructs by adding to the cells, plated in DMEM with 10% FBS, OPTI-MEM, Lipofectamine 2000, pBABE DNA, and pCL-101A packaging components. At day 1, 2, and 3, the virus-containing growth medium was removed, frozen at -80°C, and filter sterilized before using with BMDM. The parental retroviral vector pBABE was from Dr. Chris Counter (Duke University Medical School). To ectopically express SIRT4 in BMDM, freshly isolated bone marrow was added to 6-well dishes in L929-conditioned, macrophage differentiation medium containing polybrene at 5μg/mL and 0.5mL virus-containing growth medium per 6-well. Cells were spinoculated by centrifugation of the plate at 1,000 x g for 30min at room temperature. Macrophage differentiation was allowed to occur for 7 days in the presence of puromycin, to select for viral insertion. On day 4 of differentiation the growth medium was changed. After 7 days (total) of differentiation the growth medium was changed to DMEM containing 10% FBS and cells were stimulated with LPS at 100ng/mL.

Knock down of SIRT4 in RAW cells

RAW cells at ~60% confluence were transfected with SIRT4 siRNA or with negative control siRNA using Lipofectamine RNAiMAX Reagent (Life Technologies) according to the manufacturer's recommendations. Forty-eight hours later cells were stimulated with LPS at 500ng/mL. SIRT4 knock down efficiency was determined by western blotting of cell protein extracts.

Sample prep for mass spectrometry analysis

BMDM were pelleted, washed 1X with PBS, re-pelleted, and stored at -80°C until lysis. Samples were thawed on ice, resuspended in 300μL of urea lysis buffer (8M urea, 50mM Tris, pH 8.0, 40mM NaCl, and 2mM MgCl2), and disrupted with a Qiagen TissueLyzer for 30sec on a setting of 30Hz. The samples were next put through 3 cycles of freeze-thaw consisting of freezing on dry ice followed by thawing in 37°C water bath. After the last thaw, samples were sonicated using a pencil sonicator at power level 3 for 3, 5sec bursts. Samples were centrifuged at 10,000 x g for 10min at 4°C and the protein concentration of supernatants determined with a BCA assay. For each sample, 500μg protein was aliquoted into 200μl for subsequent processing steps. Samples were reduced in 10 mM dithiothreitol (VWR, VW1506-02), alkylated in 20 mM iodoacetamide (Calbiochem 407710), and digested with 200 ng of sequencing grade modified trypsin (Promega V5111) overnight in 37°C water bath. After digestion, samples were acidified to 0.5%, v/v, trifluoroacetic acid and centrifuged at 10,000 x g for 10min. Supernatants were desalted using Waters 50mg resin tC18 SEP_PAK SPE columns. Eluents were dried down and peptides were reconstituted in 200 μL of 1% TFA/2% ACN, and transferred to total recovery LC vials (Waters) for LC/MS analysis.

Mass spectrometry analysis

Label-free quantitative nanoLC-MS/MS of 2 μg of digested peptide from all experimental samples was performed using data-dependent acquisition (DDA) on a Q Exactive Plus Orbitrap mass spectrometer (ThermoFisher Scientific) coupled to an EASY-nLC UPLC system (Thermo). Samples were run in random order, interspersed by three injections of a QC pool, made from combining equal amounts of each experimental sample. For each sample, 2 μg (5 μL) of peptide was injected and tapped on an Acclaim PepMap (Thermo) trapping (3 μm, 75 μm × 20 mm) and separated on an analytical (2 μm 100 C18, 75 μm × 500 mm column) column over a 105 min gradient (5 to 40% solvent B (90% ACN/0.1% FA)) at a flow rate of 300 nL/min and a column temperature of 55°C. MS1 spectra (precursor ions) were collected at a resolution (r) of 70,000, a target AGC target value of 3e6 ions, and a maximum injection time (IT) of 100 ms. Using a DDA top20 method, precursor ions were selected using a 1.2 m/z isolation window and normalized collision energy of 27, with dynamic exclusion (DE) enabled for 30 s. MS2 spectra (product ions) were collected at r = 17,500, AGC=1e5, and max IT=100 ms. All Thermo .raw files were uploaded to Proteome Xchange, along with additional metadata on each experiment.

Proteomic data analysis

Raw data for all experimental samples and QC pools were processed in Proteome Discoverer 3.1 (PD3.1, ThermoFisher). The Minora Feature Detector node was used to align precursor ion features across all runs based on mass and retention time. Relative peptide abundance was calculated using precursor intensity, considering MS1 extracted ion chromatograms (XIC) of the aligned features. Data were searched with Sequest HT, MS Amanda 3.0, and PMI-Byonic against a mouse complete proteome database downloaded from UniProt and corresponding "decoy" reversed protein sequences. Considered modifications included oxidation (15.995 Da on M) and itaconylation (112.016 Da on K) as variable, and carbamidomethyl (57.021 Da on C) as fixed, with up to 2 missed cleavages (full trypsin specificity). FDR for peptide spectral matches (PSMs) for each search algorithm was estimated using Percolator, after which IMP-ptmRS was used to localize modifications to specific residues. Peptide Validator was used to collapse identifications to 1% FDR at the PSMs and peptide levels. Quantitation was normalized for total peptide signal within each sample, providing Normalized Abundance values to adjust for subtle differences in sample loading and LC-MS performance. Peptides were grouped to proteins (strict parsimony), which were filtered to 1% FDR with Protein FDR Validator, with only unmodified peptides that were unique to each protein group used for protein level quantitation. PD 3.1 was used to perform statistical comparisons across each genotype-treatment group, calculating Log2 fold change (Log2 FC), p-value (ANOVA), and adjusted p-value (Benjamii-Hochburg). Individual itaconylated peptides were validated using PEP scores as described in the Results section and Figure 3.

Multi-omic data integration

Multi-omics integration using GAUDI (Group Aggregation via UMAP Data Integration) was performed to analyze the relationships between metabolomic, proteomic, and itaconylated peptide datasets. Due to unequal sample sizes across the three datasets, we implemented a balancing strategy to ensure proper representation across all genotype-treatment combinations. Our custom balancing function addressed sample size discrepancies by either downsampling groups with excess samples or generating synthetic samples for underrepresented groups based on the statistical properties of existing samples. We targeted 20 total samples distributed evenly across genotype (WT/KO) and treatment (LPS 0h, LPS 6h, and LPS 24h) conditions. The balanced datasets underwent standardization with log-transformation and z-score normalization, with missing values imputed using a minimum value approach. The processed datasets were then integrated using the GAUDI method, with UMAP parameters set to n_neighbors=15 and n_components=4 for individual omics embedding, followed by n_neighbors=15 and n_components=2 for the concatenated embedding. Random Forest was selected as the feature importance method to identify key metabolites and proteins contributing to sample variance. Gene ontology enrichment analysis was performed on the integrated dataset using the org.Mm.eg.db annotation database to identify biological processes associated with the observed patterns.

BCKDH activity assay

Bone marrow from SIRT4KO mice was differentiated into BMDM on uncoated, 10cm petri dishes. Differentiated BMDM were washed 2X with PBS and then dissociated from the dishes by adding 2mL ice-cold Enzyme-Free PBS-Based Cell Dissociation Media (Gibco #13151014) and incubating dishes at 4°C for 10min. Cells were collected by pipet, pelleted by centrifugation at 1,000 x g for 10min, and plated on an XF24 Seahorse cell culture microplate. The next day the cells were washed 1X with MAS buffer (220mM mannitol, 70mM sucrose, 10mM KH2PO4, 5mM MgCl2, 2mM HEPES, 1mM EGTA, and 0.2% fatty acid free BSA) followed by the addition of 0.4mL of assay buffer to each well. Assay buffer contained 1X MAS buffer, 4mM ADP, 2mM NAD+, 0.1mM Coenzyme A, 1nM Seahorse XH Plasma Membrane Permeabilizer (PMP); Agilent, #102504, and fuel molecules. With pyruvate serving as fuel, assay buffer contained 10mM pyruvate and 0.5mM L-malate and with KIC serving as the fuel source, assay buffer contained 1mM alphaKIC and 0.3mM thiamine pyrophosphate. Following addition of assay buffer to cells the microplate plate was processed as soon as possible by the XF24 instrument. OCR measurements were normalized by protein content per well.

Wound healing assay

Mice were anesthetized in a holding box with 4% isoflurane and then moved to nose cones at which point the isoflurane was reduced to 2%. Ophthalmic lube was applied to the eyes and buprenorphine HCl, at 0.05-0.1mg/kg, was pre-operatively injected subcutaneously. Clippers were used to remove hair from the upper back in the area to be wounded. The surgical area on the animals was wiped alternatingly with 70% ethanol and providone-iodine, 3X. Before generating wounds, the animals were injected subcutaneously again, this time with 1mg/kg buprenorphine Lab SR (supplied at 1mg/kg). Excisional wounds were then generated using 5mm biopsy punches while holding skin away from the middorsal line. Wound diameter was measured with a digital caliper and the animals were allowed to recover on a heating pad. Once animals were ambulatory and alert they were transferred to clean recovery cages. Male and female mice (n=3 per genotype per sex) were used for wound healing studies, with measurements taken over 9 days.

QUANTIFICATION AND STATISTICAL ANALYSIS

Data are presented as mean ± SEM unless otherwise indicated. Statistical analyses were performed using R (version 4.0) or GraphPad Prism software (version 9.0). All code for statistical analyses is publicly available (https://github.com/hirscheylab/itaconylation?tab=readme-ov-file) For Prism analyses, differences between two groups were analyzed using unpaired two-tailed Student's t-test. Differences among multiple groups were analyzed by one-way or two-way ANOVA followed by appropriate post-hoc tests. For wound healing experiments, two-way repeated measures ANOVA was used to analyze differences between genotypes over time. Statistical significance was set at p < 0.05. Sample sizes for each experiment are indicated in the figure legends. For proteomics data, statistical significance was determined using ANOVA with Benjamini-Hochberg correction for multiple comparisons. A threshold of adjusted p < 0.05 and absolute log2 fold change > 0.5 was used to define significantly changing proteins or peptides.

ADDITIONAL RESOURCES

Raw metabolomics and immune data are deposited at <https://zenodo.org/records/15372985>

Raw proteomics are deposited at XYZ
