## Supplementary material for "SIRT4 Controls Macrophage Function and Wound Healing through Control of Protein Itaconylation in Mice": Reagents

**Key resources table**

| REAGENT or RESOURCE | SOURCE | IDENTIFIER |
| --- | --- | --- |
| Antibodies | | |
| Mouse monoclonal anti-IRG1 | Cell Signaling Technology | 77510 |
| Mouse monoclonal anti-OxPhos mix | Abcam | ab110413 |
| Rabbit polyclonal anti-SIRT4 | Sigma-Aldridge | HPA029691 |
| Rabbit polyclonal anti-itaconyl lysine polyclonal | This paper |  |
| Rabbit polyclonal anti-succinyl lysine | This paper |  |
| Bacterial and virus strains | | |
| Bacteria: BL21 (DE3) pLysS | Thermo Fisher Scientific | C606003 |
| Biological samples |  |  |
| Mouse serum | This paper | N/A |
| Mouse bone marrow | This paper | N/A |
| Chemicals, peptides, and recombinant proteins | | |
| Lipopolysaccharide (LPS) | Invitrogen | co-4976-93 |
| Bovine serum albumin, fatty acid free | Sigma-Aldridge | A7030 |
| Itaconic anhydride | Sigma-Aldridge | 259926 |
| Acetic anhydride | Sigma-Aldridge | 91204 |
| 3-Methyl-glutaric anhydride | Sigma-Aldridge | M47809 |
| Glutaric anhydride | Sigma-Aldridge | G3806 |
| Acetylated BSA | This paper | N/A |
| Methyl-glutarylated BSA | This paper | N/A |
| Glutarylated BSA | This paper | N/A |
| Itaconylated BSA | This paper | N/A |
| 32P-nicotinamide adenine nucleotide | Perkin Elmer | NEG023 |
| HALT protease inhibitor cocktail | Thermo Fisher Scientific | 78420B |
| SIRT4 recombinant protein | This paper | N/A |
| Enzyme-Free PBS-Based Cell Dissociation Media | Gibco | 13151014 |
| AKC Lysing Buffer | Gibco | A1049201 |
| Plasma Membrane Permeabilizer (PMP) | Agilent | 102504-100 |
| Critical commercial assays | | |
| Mouse IL-1 beta/IL-1F2 DuoSet ELISA kit | R&D Systems | DY401-05 |
| V-PLEX Mouse Cytokine 19-plex Kit | Meso Scale Diagnostics, LLC | K15255D |
| Lipofectamine transfection reagent RNAiMAX | Thermo Fisher Scientific | 13778100 |
| Deposited data | | |
| Mass spectrometry proteomics data | This paper | ProteomeXchange: PXD063862 |
| jPOST | This paper | JPST003806 |
| Experimental models: Cell lines | | |
| Mouse: RAW 264.7 | Laboratory of Anthony Means |  |
| Mouse: L929 cell line | Laboratory of Greg Taylor |  |
| Human: 293T cell line | ATCC | CRL-3216 |
| Experimental models: Organisms/strains | | |
| Mouse: Whole-body SIRT4KO: B6.129-Sirt4tm1.1Mcby/J | Jackson Laboratory | Stock#012756 |
| Mouse: C57BL/6J | Jackson Laboratory | Stock#000664 |
| Mouse: C57BL/6NJ | Jackson Laboratory | Stock#005304 |
| Oligonucleotides | | |
| Sirt4 F primer: (5’-GTCTGTCCTAGCTTCCTCACTG-3’) | Custom-made by Integrated DNA Technologies | N/A |
| Sirt4 R1 primer: (5’-TAAAGATAGTTGTAAGTCACC-3’) | Custom-made by Integrated DNA Technologies | N/A |
| Sirt4 R2 primer: (5’-AGAGCCCAGTGTGCTGGGTTG-3’) | Custom-made by Integrated DNA Technologies | N/A |
| Mouse SIRT4 siRNA | Santa Cruz Biotechnologies | sc-63025 |
| Control siRNA-A | Santa Cruz Biotechnologies | sc-37007 |
| Recombinant DNA | | |
| GST-mSIRT4(24-333) | This paper | N/A |
| pGEX-6P1 | GE Healthcare | #28-9546-48 |
| pBABE | The laboratory of Chris Counter | N/A |
| pBABE-mSIRT4 | This paper | N/A |
| pCL-101A packaging components | The laboratory of Chris Counter | N/A |
| Software and algorithms | | |
| Proteome Discoverer 3.1 | Thermo Fisher | N/A |
| LI-COR Odyssey Infrared Imaging System software | LI-COR | N/A |
| GAUDI (Group Aggregation via UMAP Data Integration) | https://github.com/pcastellanoescuder/GAUDI | N/A |
| Other | | |
| XF24 Seahorse Analyzer | Agilent | N/AA |
| LI-COR Odyssey Infrared Imaging System | LI-COR Biosciences | N/A |
| Silica-gel coated thin layer chromatography plate | Sigma-Aldrich | 60768 |
| STORM820 phosphorimager | GE Healthcare | N/A |
| Q Exactive Plus Orbitrap mass spectrometer | Thermo Fisher Scientific | N/A |
| EASY-nLC UPLC system | Thermo Fisher Scientific | N/A |
| Biopsy punch (5mm) | Medical Device Depot | 33-35 |
