## Supplementary material for "SIRT4 Controls Macrophage Function and Wound Healing through Control of Protein Itaconylation in Mice": Uncropped blots

### Slide 1
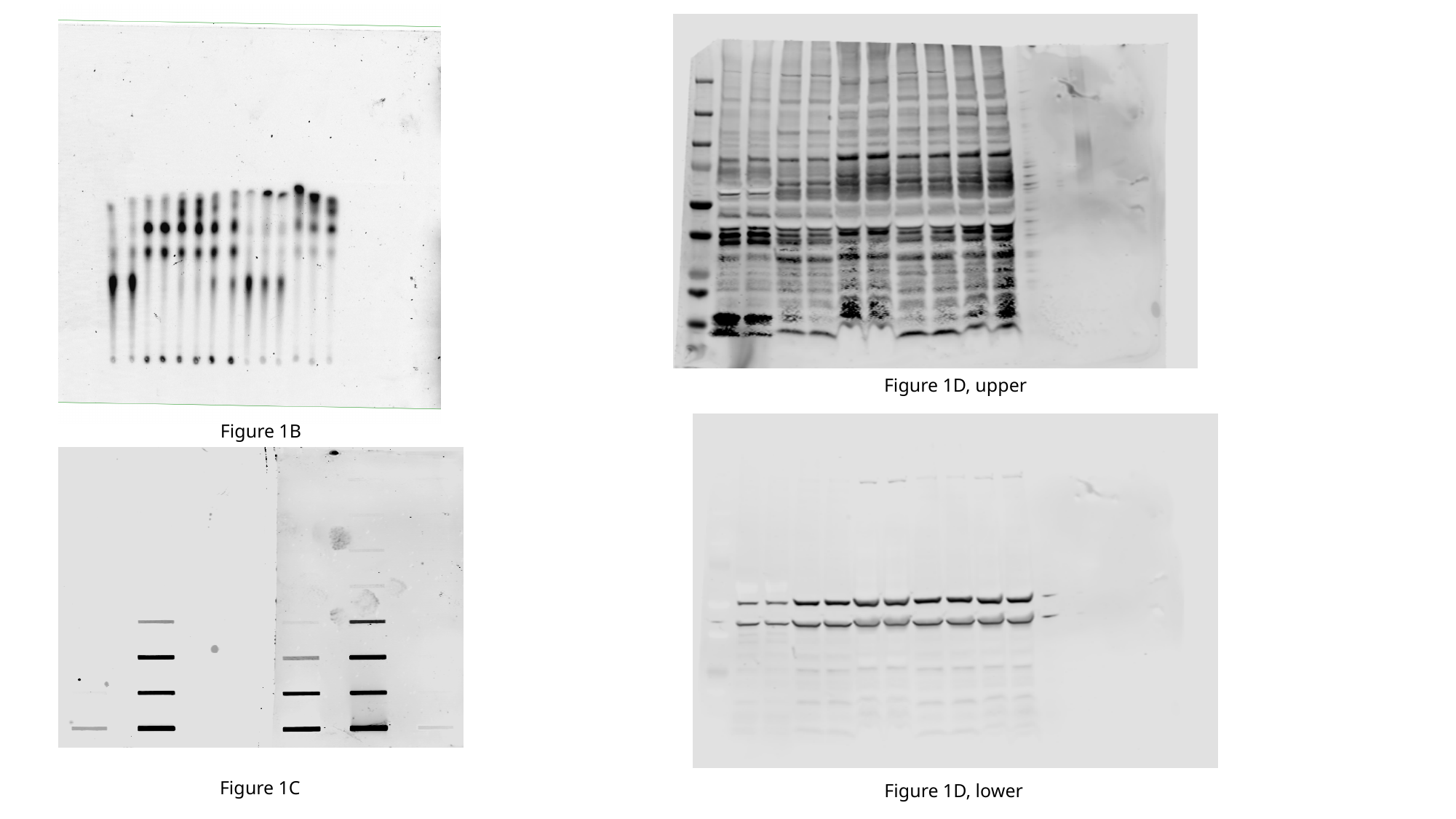

Figure 1D, upper
Figure 1B
Figure 1C
Figure 1D, lower

### Slide 2
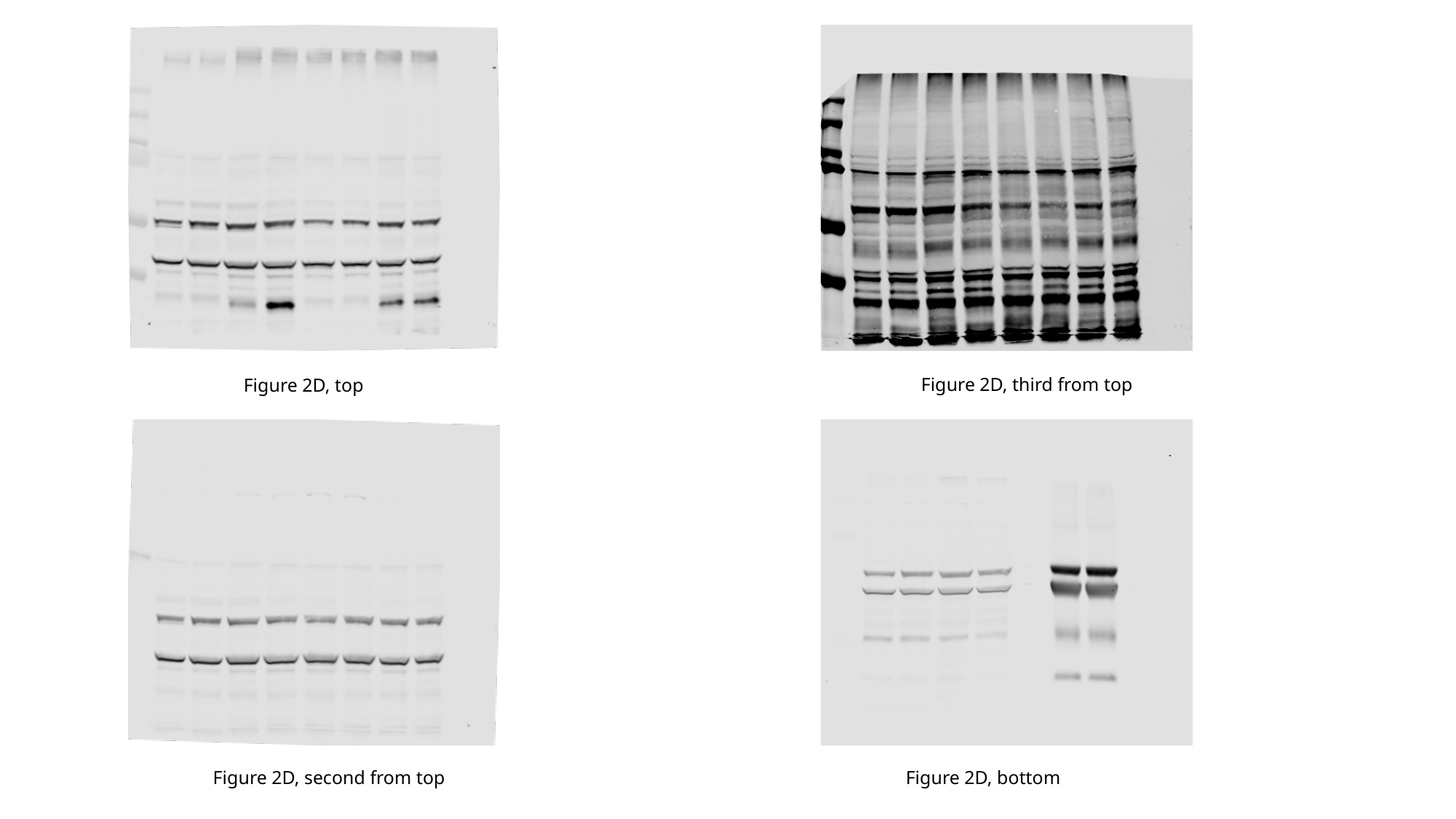

Figure 2D, third from top
Figure 2D, top
Figure 2D, second from top
Figure 2D, bottom

### Slide 3
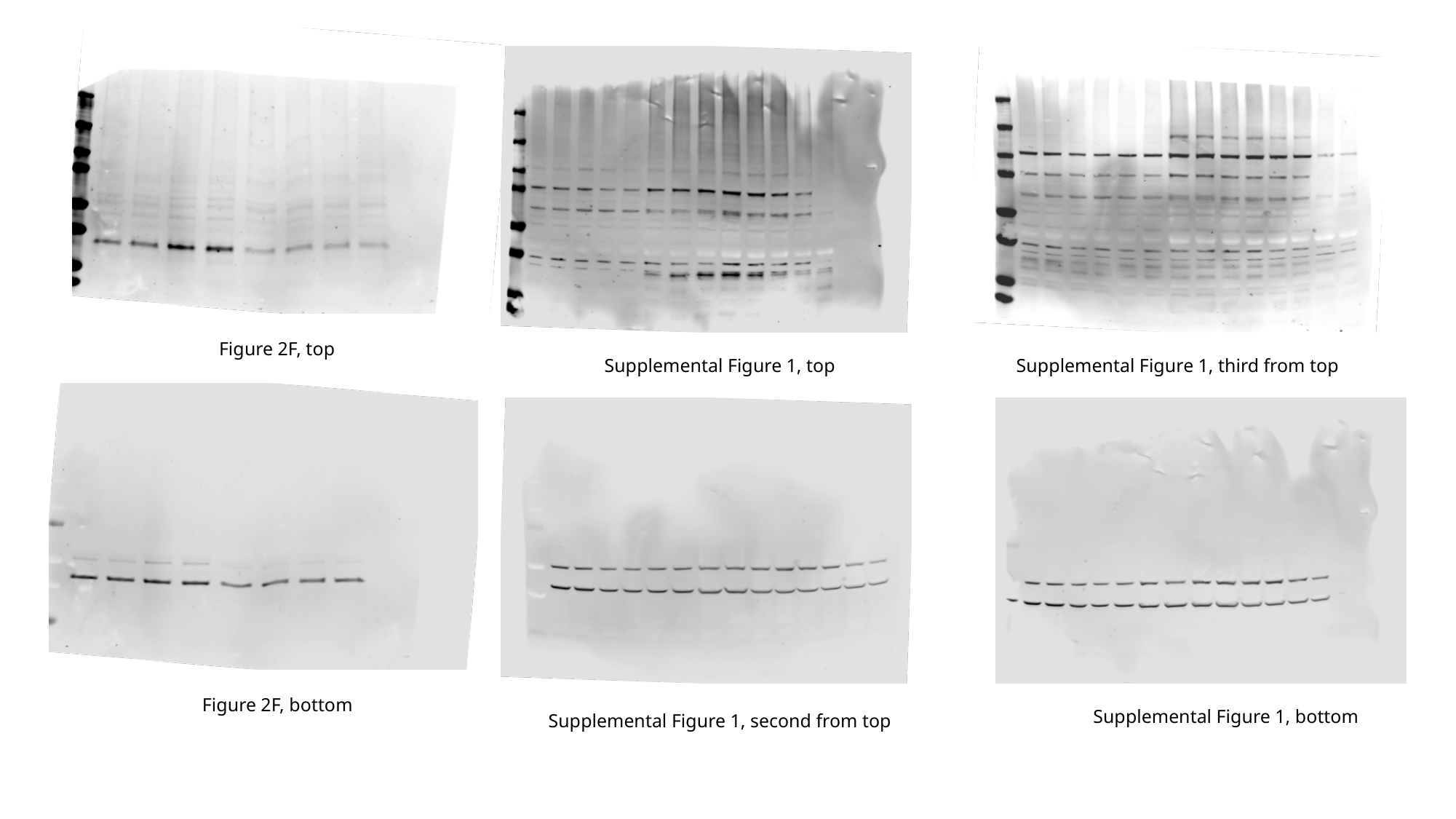

Figure 2F, top
Supplemental Figure 1, third from top
Supplemental Figure 1, top
Figure 2F, bottom
Supplemental Figure 1, bottom
Supplemental Figure 1, second from top
